# IDSL.GOA: Gene Ontology Analysis for Interpreting Metabolomic datasets

**DOI:** 10.1101/2023.03.25.534225

**Authors:** Priyanka Mahajan, Oliver Fiehn, Dinesh Barupal

## Abstract

Biological interpretation of metabolomic datasets often ends at a pathway analysis step to find the over-represented metabolic pathways in the list of statistically significant metabolites. However, definitions of biochemical pathways and metabolite coverage vary among different curated databases, leading to missed interpretations. For the lists of genes, transcripts and proteins, Gene Ontology (GO) terms over-presentation analysis has become a standardized approach for biological interpretation. But, GO analysis has not been achieved for metabolomic datasets. We present a new knowledgebase (KB) and the online tool, Gene Ontology Analysis by the Integrated Data Science Laboratory for Metabolomics and Exposomics (IDSL.GOA) to conduct GO over-representation analysis for a metabolite list. The IDSL.GOA KB covers 2,393 metabolic GO terms and associated 3,144 genes, 1,492 EC annotations, and 2,621 metabolites. IDSL.GOA analysis of a case study of older vs young female brain cortex metabolome highlighted 82 GO terms being significantly overrepresented (FDR <0.05). We showed how IDSL.GOA identified key and relevant GO metabolic processes that were not yet covered in other pathway databases. Overall, we suggest that interpretation of metabolite lists should not be limited to only pathway maps and can also leverage GO terms as well. IDSL.GOA provides a useful tool for this purpose, allowing for a more comprehensive and accurate analysis of metabolite pathway data. IDSL.GOA tool can be accessed at https://goa.idsl.me/

## Introduction

Metabolomics enables the simultaneous study of multiple metabolic processes, including pathways, transport, and reactions. Metabolomics assays are diverse and complex in terms of their analytical conditions, but they can generate quantitative and semi-quantitative data for hundreds of endogenous metabolites^1^. Recently reported datasets can have between 1,500 to 2,000 named metabolites and several thousand unidentified metabolites^1, 2^. These metabolites originate from overlapping pathways of catabolic and anabolic reactions and can also be biomarkers for metabolic processes^3^. Environmental, genetic, or biological factors can alter the regulatory, signaling, and enzyme kinetic mechanisms in one or more metabolic pathways and processes, leading to altered levels of related metabolites in cells, tissues or body fluids^4, 5^. For example, aging reprograms carbohydrate and lipid metabolism pathways in the liver^6^, tobacco smoke exposure alters the nucleotide and reactive oxidative stress species metabolism^7^, and FADS gene polymorphisms alter the levels of circulating PUFAs^8^. We can expect to see a continuous growth in the number of named metabolites in metabolomic datasets due to new advances^9, 10^ in analytical techniques and computational methods and resources.

One of the key challenges in utilizing metabolomic datasets is how to interpret these large chemical lists for mechanistic insights^11^. Pathway and network analysis can provide mechanistic insights into the biological pathways linked to the altered metabolites^12^. Interestingly, metabolomic datasets often have metabolites that are yet to be connected to a biochemical reaction and pathway^13, 14^. To also include these poorly studied metabolites, hybrid approaches of the atomic mapping of reaction and chemical similarity network (MetaMapp) and enrichment analysis (ChemRICH) can be used^13, 14^. Transcripts and protein lists are also often interpreted using gene ontology (GO) term enrichment analysis^15^, which covers terms that relate to pathways as well as other biological processes such as cell cycle or apoptosis, or even pathways that are not yet included in other biochemical databases. However, there is not yet a single tool developed that can perform a GO analysis for a metabolite list.

We have developed a new tool named ‘IDSL.GOA’ (Gene Ontology Analysis by the Integrated Data Science Laboratory for Metabolomics and Exposomics) to perform GO enrichment analysis for a list of metabolites. The tool is supported by a knowledge base of genes, enzymes, and reactants (metabolites) that are directly sources from National Center for Biotechnology Information (NCBI), Expasy and GO consortium databases. We present a case study of an aging mouse metabolic atlas to highlight the metabolic processes that were suggested to be related to the aging process and were only identified by the IDSL.GOA based GO analysis method. The online tool is available at https://goa.idsl.me/ site.

## Material and methods

*IDSL*.*GOA Knowledgebase:* We assembled and integrated information from a diverse set of data sources, including genes, enzymes, compounds, gene ontology terms and the relationships among them. Table 1 provides the web addresses for the publicly available data sources and their respective locations. To focus specifically on metabolism, we restricted our gene selection to those related to GO term GO:0008152 (metabolic process) and linked with the human genome. Only the downstream entities for these metabolic genes were included in the knowledgebase. Utilized identifiers for creating the knowledgebase were - NCBI Gene, NCBI Protein, NCBI Nucleotide, GO Term, Enzyme Commission Number (EC) and InChiKeys. Linkages among these entities were extracted or accessed from the resources listed in Table 1.

**Table 1.** Data sources for assembling the IDSL.GOA knowledgebase.

| Resource | Address | Information |
| --- | --- | --- |
| Gene Ontology | http://purl.obolibrary.org/obo/go.obo | GO-GO |
| Expasy Enzyme | https://ftp.expasy.org/databases/enzyme/enzyme.dat | EC Numbers |
| NCBI Gene Accession | https://ftp.ncbi.nlm.nih.gov/gene/DATA/gene2accession.gz | Gene – Transcript<br>Transcript - Protein |
| NCBI Gene GO Annotation | https://ftp.ncbi.nlm.nih.gov/gene/DATA/gene2go.gz | Gene - GO |
| NCBI PubChem | https://pubchem.ncbi.nlm.nih.gov/ | EC-compounds |

Note: These information were accessed in December 2023.

*Over-representation statistics:* For the GO analysis, we employed an overrepresentation analysis (ORA) test using the hypergeometric distribution. This statistical test is a widely accepted method for determining whether a set of molecular entities (gene or proteins or metabolites) is significantly overrepresented in a particular biological pathway or process, given a background database. We also applied filters 1) overlap >= 3, 2) at least three genes in the GO process 3) The set size < 5% of total compounds 4) FDR < 0.05 and 5) the overlap should be >5% of the total set size for a GO term. The overlap represents how many out of the input InChiKey list are found among the InChiKey identifiers linked with a GO term. Only the first 14 characters of an InChiKey, which represent the two-dimensional structure were used to find the overlap. These filters narrow down the list of GO terms to only the most relevant ones.

We have used “phyper” function in R to compute the hypergeometric test. The parameter for the test were – *phyper(x-1,y,a,b, lower*.*tail = FALSE)*, where x is the overlap between the input list of InChiKey and compounds linked with a GO term, *y* is the count of all compounds(2D structures) linked with the GO term, *a* is the count of all compounds (2D structures) not linked with the GO-term (1,856-y), *b* is the count of the InChiKey from the input list that were found in the knowledgebase. By default, the *phyper* function in R calculates the probability of drawing less than or equal to *x* for a GO term. Use of the parameters “x-1” and “lower.tail=FALSE” returns the probability of drawing more than or equal to *x* for a GO term. The total number of compounds (2D structures) linked with GO terms was 1,856. For example, for the Nucleoside salvage (GO:0043174), *x* was 12, *y* was 58, *b* was 73, and *a* was 1,798 (1,856-58) for the test study’s results. The *p*-value of this GO term was computed as ‘*phyper(11,58,1784,73, lower*.*tail = FALSE)’* which returns 1.158242e-06.

The IDSL.GOA tool uses the False Discovery Rate (FDR) cutoff of 0.05 to control the proportion of false positives in multiple hypothesis testing in GO analysis. We repeated this test for all metabolically relevant 2,392 GO-terms.

*Case study and its analysis*. Our test study was based on publicly available data from the Aging Mouse Brain Metabolome Atlas^1^, a comprehensive resource that provides information on the metabolites found in the different regions of brain of aging mice. Specifically, we compared the brain metabolome of the cortex region in an older female mouse against that of a young mouse. To identify the significantly different metabolites, we used the student t-test. We used InChiKey identifiers for the compounds that had a *p*-value of less than 0.05 in the student t-test.

*IDSL*.*GOA online tool:* The online tool was developed using the ReactJS JavaScript framework (https://reactjs.org/), which is known for its efficient rendering of dynamic user interfaces. To facilitate data visualization, we utilized the Google Chart (https://developers.google.com/chart) and Cytoscape JS plugins (https://github.com/plotly/react-cytoscapejs), specifically designed to work with ReactJS. Cytoscape online version is a lightweight and user-friendly tool that allows users to perform basic network visualization and analysis tasks without the need to install the software locally. For small networks, the online version may be sufficient, but for larger and complex network, it is recommended to download the Cytoscape SIF (Simple Interaction Format) file and use the local version of Cytoscape software to create high resolution graphics. Instructions to use the IDSL.GOA tool are provided on the landing page.

## Results

### Creating the IDSL.GOA metabolic knowledgebase

To perform IDSL.GOA over-representation analysis, we first needed to create a database of relationships among metabolic entities. This database was designed to capture the heterogenous relationships among genes, enzymes, compounds, and gene ontology terms. The source data for these relationships were obtained from various publicly available key databases, including the NCBI, Expasy – SIB Swiss Institute of Bioinformatics, and the Gene Ontology Consortium (Table 1). We restricted the knowledgebase to only human genes and their products in the first version of the KB. The resulting version 1 of the IDSL.GOA database contained a total of 3,144 genes, 1,492 enzyme commission numbers, 2,621 compounds, 1,856 2D chemical structures and 2,393 gene ontology terms for metabolic processes (Figure 1). Overall, the IDSL.GOA database provided a comprehensive resource for performing GO over-representation analysis for metabolite lists.

**Figure 1.**
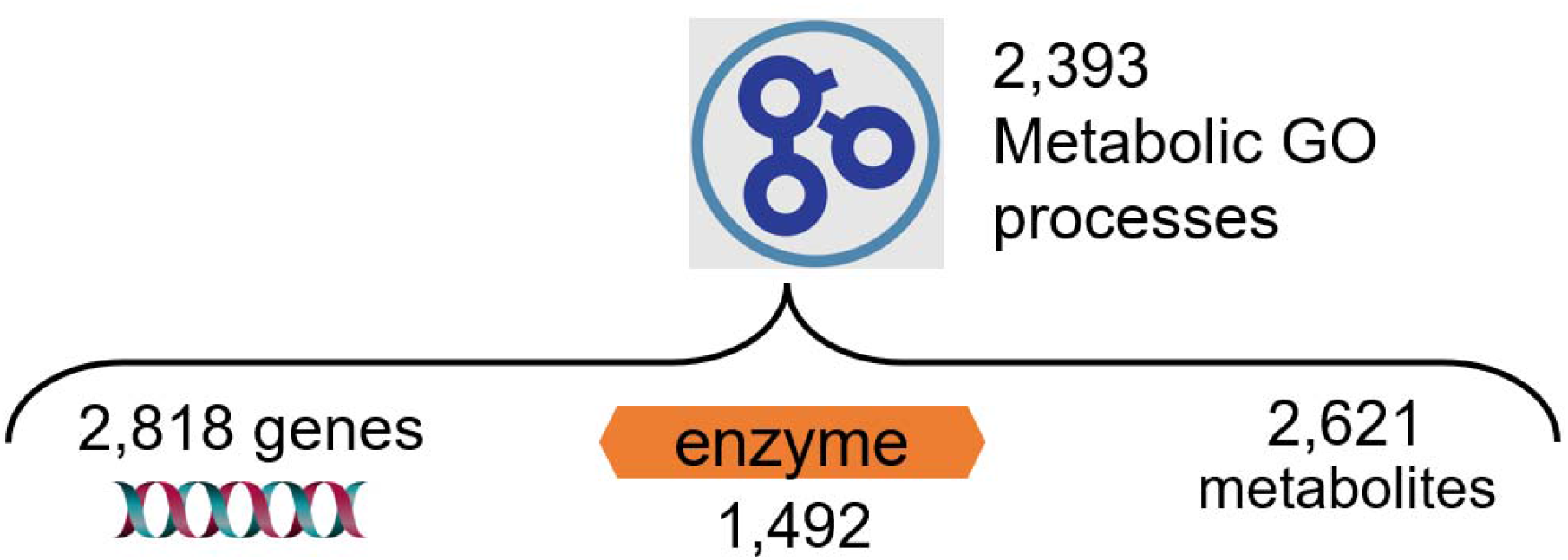
Content and relationships in the IDSL.GOA metabolic knowledgebase. Total number of metabolic GO terms under the metabolic process (GO:0008152) are 6,084.

### Aging mouse brain metabolomics-a case study

In this study, we aimed to investigate the changes in metabolite levels in the brain cortex of old and young mice using a metabolomic atlas that contained close to 1,547 identified compounds. We identified 557 metabolites that were significantly different between the old (59 weeks) and young (3 weeks) female mouse brain cortex (Table S1). InChiKeys for these significant metabolites were used as input for IDSL.GOA analysis. The GO analysis results suggested a total of 82 GO processes that were over-represented in the input list at an FDR cutoff of 0.05 (Table S2). The GO network and the impact plot visualization suggested that processes in nucleotide and amino acid metabolism (GO:0043174, GO:0046415 and GO:0006166) were significantly affected during the aging process (Figure 2-3, Table S2, Figure S1).

**Figure 2.**
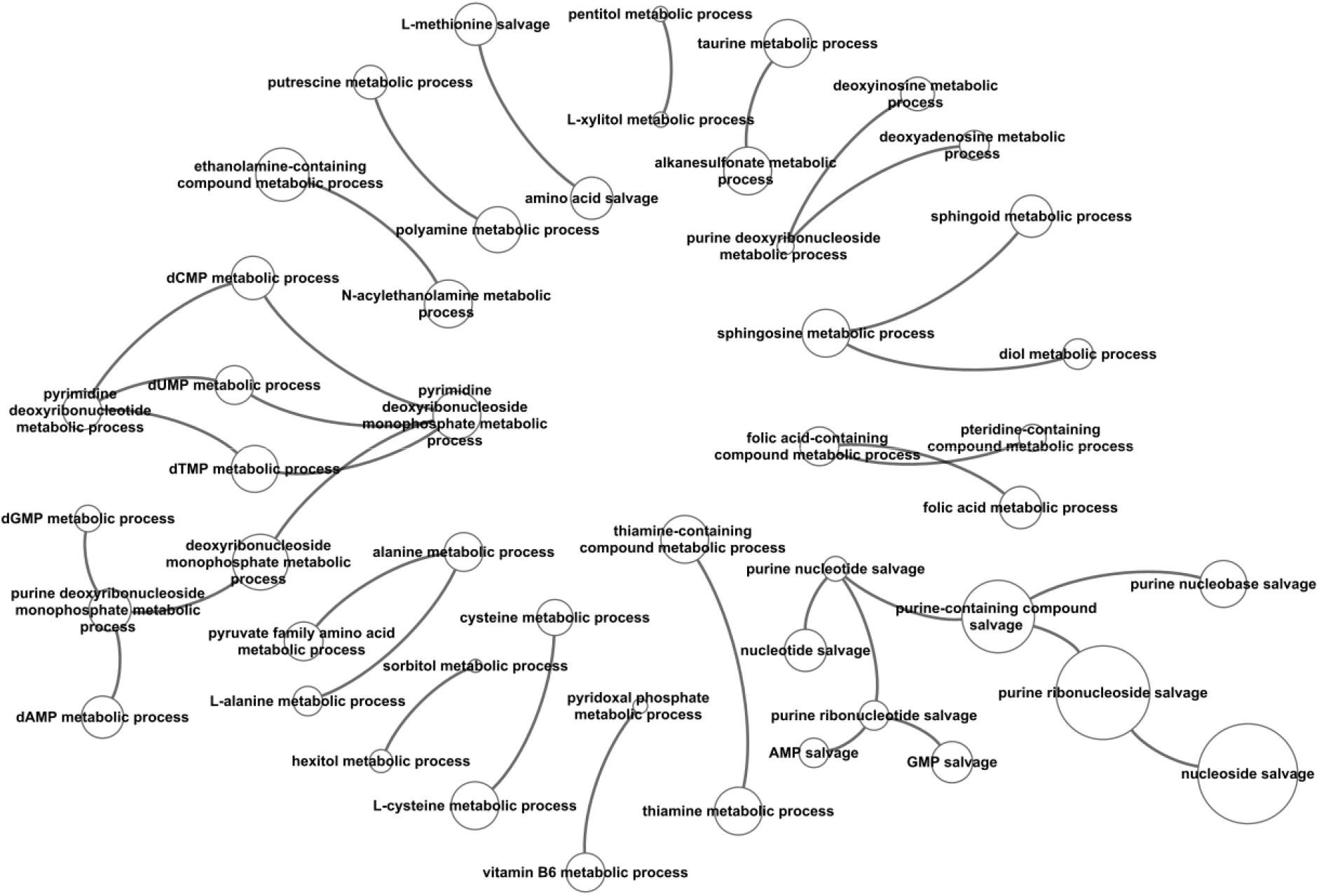
GO Tree visualization of the significantly overrepresented GO-terms in the input metabolite list. Node size indicates the statistical significance, larger ones are most significant.

**Figure 3.**
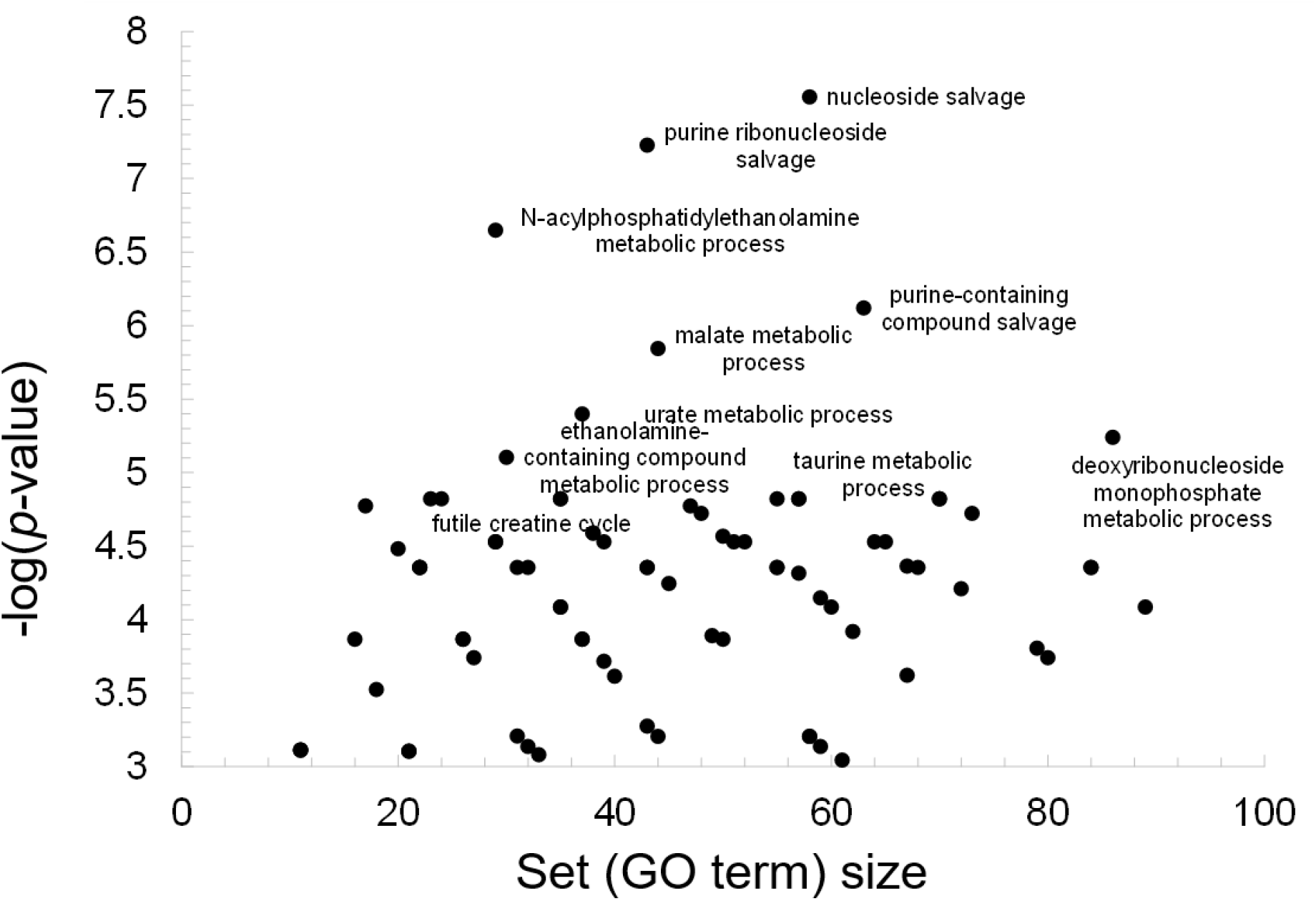
IDSL.GOA impact plot to show the most overrepresented GO terms by their specificity. A small set size shows more specific metabolic processes. For clarity, only the top metabolic processes are labelled but table S2 has the names for all the significant metabolic processes.

### IDSL.GOA online tool

The IDSL.GOA online tool is a user-friendly resource for identifying overrepresented metabolic processes in a list of metabolites. The online interface offers features including analysis, query, explore, statistic and download options. The ‘Run Go Analysis’ option on the landing page allows users to input a list of InChiKeys and obtain results in various formats, including Cytoscape SIF, Microsoft Excel, and CSV. The InChiKeys for only the significant compounds (p<0.05) in a statistical test should be used as input. The Cytoscape SIF and node attribute files are useful for creating high-resolution figures in the Cytoscape desktop software^16^. The primary analysis results are visualized in a ‘GO Ontology network’ graph using Cytoscape JS library, which provides an intuitive and interactive way to explore the data. This view is analogous to the pathway ontology visualization in the Reactome database^17^. The size of the node in the graph reflects the significance of the term, with larger nodes indicating more significant terms in the hypergeometric test. Additionally, an impact plot shows how specific the GO terms are for the input list, by plotting the set size versus -log(p-value). The explore option allows users to navigate the GO ontology tree. Clicking on a GO term in the main analysis, query or explore options provide the GO-term specific InChikeys that overlap with the input list. The query option allows users to query a single compound, reaction, gene, protein and transcript to retrieve the associated metabolic GO terms. All GO network visualization has a basic set of layouts (views) implemented which can be explored by a user to find the most readable and helpful views for a GO ontology network that can aid in the biological interpretation of metabolite lists. Finally, the statistics and download tabs provide updates on the database version and download links, and the landing page offers Instructions for using the database.

## Discussion

IDSL.GOA is the first bioinformatics tool that used GO terms for over-representation analysis of metabolomic datasets. By mapping the metabolites to their associated GO terms, IDSL.GOA can improve the mechanistic interpretation of metabolomics data by providing a functional annotation of the metabolites based on their associated metabolic processes and pathways in the Gene Ontology database. It is a more sensitive and accurate tool for data with larger lists (>1,000 named metabolites)^1, 2^. This can lead to the identification of key regulatory pathways and molecular mechanisms that are involved in the observed changes and can guide further experimentation and hypothesis testing. By leveraging the new IDSL.GOA knowledgebase, we were able to identify the overrepresented metabolic pathways and processes in our case study dataset and gain new insights into the underlying mechanisms that govern metabolic activity in aging brain tissue.

### Advantages of using GO terms for metabolomics data interpretation

There are several advantages of GO analysis over traditional pathway analysis. GO analysis provides a more comprehensive annotation system for genes and their products than pathway analysis, allowing for a broader range of metabolic processes and pathways to be analyzed^18^. Unlike pathway analysis, GO analysis is not limited to hand-drawn pathway maps which tend to differ from one database to another, making it more flexible and adaptable to different experimental conditions. Depending on the background pathway database, the interpretation of metabolite lists can differ and may be inaccurate, leading to contradicting results and less impact^3^. On contrast, GO analysis allows for a more detailed and accurate interpretation of results, as it provides a broader context for the function and regulation of metabolite levels. Because the GO system is standardized, it allows for greater consistency and comparability between different studies and datasets. GO terms not only covers the known pathway maps but also covers additional metabolic processes that are not yet included in the pathway databases.

### Key strengths of IDSL.GOA tool

The IDSL.GOA tool is a free, user-friendly and web-based platform that utilizes Gene Ontology (GO) terms for the analysis of metabolomics data. It offers an intuitive interface that allows users to perform GO enrichment analysis for an input metabolite list. The tool has a range of useful features to facilitate the interpretation and has a wide range of capabilities, including query, explore, statistics, and download options. The use of GO terms provides an improved biological interpretation of metabolomics data, which can help researchers identify novel and metabolically relevant pathways and processes. The tool is built on a robust knowledgebase that contains relationships among metabolic entities, obtained from various sources including NCBI, Expasy and the Gene Ontology Consortium databases. The tool allows for a more comprehensive and accurate analysis of metabolomics data by identifying not only the predefined pathways but also relevant metabolic processes that are not included in the commonly used pathway databases. It is the first of its kind tool for metabolomics data.

### Future plan

In the follow up work, we plan to improve IDSL.GOA by expanding the underlying database of GO to metabolite relationships. For this, we will curate and map the annotated compounds in the publicly available metabolomic datasets to the enzyme activity annotations and subsequently to the GO terms. There is also a need to harmonize the chemical information across reaction databases and metabolomics reports. Since the mapping between GO terms to genes, proteins and transcripts is available, the future version of IDSL.GOA may also facilitate a multi-omics GO analysis.

### Limitations

Few limitations should be noted. The IDSL.GOA tool relies on the availability of InChiKey-linked metabolite data, and the coverage of metabolite curation may vary across different metabolomics laboratories. Not all annotated compounds in the metabolomic datasets have been linked with EC numbers in the biochemical databases. The GO hierarchy and associated annotations may contain biases or inaccuracies due to incomplete or outdated information. There is some redundancy in GO term names which may inflate the over-representation analysis results. The mechanistic interpretation still needs to be validated by additional experimentation. By discussing these limitations, we can provide a more balanced view of the capabilities and potential drawbacks of the IDSL.GOA tool for GO analysis in metabolomics.

## Conclusions

In summary, the IDSL.GOA tool can enable a comprehensive and accurate biological interpretation of metabolomics data. A much-needed transition from pathway maps to GO terms for interpreting metabolomic datasets can be supported by the IDSL.GOA tool. It is more sensitive in identifying significantly enriched GO terms that are relevant for metabolic processes. By providing a comprehensive view of the underlying biology, this approach can facilitate the identification of key regulatory pathways and biomarkers that may be useful for diagnosis, prognosis, and therapeutic targeting.

## Tables Funding

NIH (U24ES035386, U2CES026561, R01ES032831, R35ES030435, U2CES026555 P30ES023515, K12ES033594, U2CES030859, UL1TR004419)

## Data availability

The Aging Mouse Metabolome Atlas dataset can be accessed at https://doi.org/10.21228/M8C68D IDSL.GOA knowledgebase elements and relationships are available at https://zenodo.org/records/10223649

## Software availability

IDSL.GOA tool can be accessed at https://goa.idsl.me/ site and https://github.com/idslme/IDSL.GOA

## References

(1) Ding, J.; Ji, J.; Rabow, Z.; Shen, T.; Folz, J.; Brydges, C. R.; Fan, S.; Lu, X.; Mehta, S.; Showalter, M. R.; et al. A metabolome atlas of the aging mouse brain. Nat Commun 2021, 12 (1), 6021. DOI: 10.1038/s41467-021-26310-y

(2) Byeon, S. K.; Madugundu, A. K.; Garapati, K.; Ramarajan, M. G.; Saraswat, M.; Kumar, M. P.; Hughes, T.; Shah, R.; Patnaik, M. M.; Chia, N.; et al. Development of a multiomics model for identification of predictive biomarkers for COVID-19 severity: a retrospective cohort study. Lancet Digit Health 2022, 4 (9), e632–e645. DOI: 10.1016/S2589-7500(22)00112-1

(3) Wieder, C.; Frainay, C.; Poupin, N.; Rodriguez-Mier, P.; Vinson, F.; Cooke, J.; Lai, R. P.; Bundy, J. G.; Jourdan, F.; Ebbels, T. Pathway analysis in metabolomics: Recommendations for the use of over-representation analysis. PLoS Comput Biol 2021, 17 (9), e1009105. DOI: 10.1371/journal.pcbi.1009105

(4) Sarkar, A.; Jin, Y.; DeFelice, B. C.; Logan, C. Y.; Yang, Y.; Anbarchian, T.; Wu, P.; Morri, M.; Neff, N. F.; Nguyen, H.; et al. Intermittent fasting induces rapid hepatocyte proliferation to restore the hepatostat in the mouse liver. Elife 2023, 12. DOI: 10.7554/eLife.82311

(5) Tanes, C.; Bittinger, K.; Gao, Y.; Friedman, E. S.; Nessel, L.; Paladhi, U. R.; Chau, L.; Panfen, E.; Fischbach, M. A.; Braun, J.; et al. Role of dietary fiber in the recovery of the human gut microbiome and its metabolome. Cell Host Microbe 2021, 29 (3), 394–407 e395. DOI: 10.1016/j.chom.2020.12.012

(6) Hunt, N. J.; Kang, S. W. S.; Lockwood, G. P.; Le Couteur, D. G.; Cogger, V. C. Hallmarks of Aging in the Liver. Comput Struct Biotechnol J 2019, 17, 1151–1161. DOI: 10.1016/j.csbj.2019.07.021

(7) Yuan, J. M.; Gao, Y. T.; Murphy, S. E.; Carmella, S. G.; Wang, R.; Zhong, Y.; Moy, K. A.; Davis, A. B.; Tao, L.; Chen, M.; et al. Urinary levels of cigarette smoke constituent metabolites are prospectively associated with lung cancer development in smokers. Cancer Res 2011, 71 (21), 6749–6757. DOI: 10.1158/0008-5472.CAN-11-0209

(8) Surendran, P.; Stewart, I. D.; Au Yeung, V. P. W.; Pietzner, M.; Raffler, J.; Worheide, M. A.; Li, C.; Smith, R. F.; Wittemans, L. B. L.; Bomba, L.; et al. Rare and common genetic determinants of metabolic individuality and their effects on human health. Nat Med 2022, 28 (11), 2321–2332. DOI: 10.1038/s41591-022-02046-0

(9) Vasilopoulou, C. G.; Sulek, K.; Brunner, A. D.; Meitei, N. S.; Schweiger-Hufnagel, U.; Meyer, S. W.; Barsch, A.; Mann, M.; Meier, F. Trapped ion mobility spectrometry and PASEF enable in-depth lipidomics from minimal sample amounts. Nat Commun 2020, 11 (1), 331. DOI: 10.1038/s41467-019-14044-x

(10) Koopman, J.; Grimme, S. From QCEIMS to QCxMS: A Tool to Routinely Calculate CID Mass Spectra Using Molecular Dynamics. J Am Soc Mass Spectrom 2021, 32 (7), 1735–1751. DOI: 10.1021/jasms.1c00098

(11) Barupal, D. K.; Fan, S.; Fiehn, O. Integrating bioinformatics approaches for a comprehensive interpretation of metabolomic datasets. Curr Opin Biotechnol 2018, 54, 1–9. DOI: 10.1016/j.copbio.2018.01.010

(12) Lind, L.; Fall, T.; Arnlov, J.; Elmstahl, S.; Sundstrom, J. Large-Scale Metabolomics and the Incidence of Cardiovascular Disease. J Am Heart Assoc 2023, 12 (2), e026885. DOI: 10.1161/JAHA.122.026885

(13) Barupal, D. K.; Fiehn, O. Chemical Similarity Enrichment Analysis (ChemRICH) as alternative to biochemical pathway mapping for metabolomic datasets. Sci Rep 2017, 7 (1), 14567. DOI: 10.1038/s41598-017-15231-w

(14) Barupal, D. K.; Haldiya, P. K.; Wohlgemuth, G.; Kind, T.; Kothari, S. L.; Pinkerton, K. E.; Fiehn, O. MetaMapp: mapping and visualizing metabolomic data by integrating information from biochemical pathways and chemical and mass spectral similarity. BMC Bioinformatics 2012, 13, 99. DOI: 10.1186/1471-2105-13-99

(15) Gene Ontology, C. The Gene Ontology resource: enriching a GOld mine. Nucleic Acids Res 2021, 49 (D1), D325–D334. DOI: 10.1093/nar/gkaa1113

(16) Shannon, P.; Markiel, A.; Ozier, O.; Baliga, N. S.; Wang, J. T.; Ramage, D.; Amin, N.; Schwikowski, B.; Ideker, T. Cytoscape: a software environment for integrated models of biomolecular interaction networks. Genome Res 2003, 13 (11), 2498–2504. DOI: 10.1101/gr.1239303

(17) Gillespie, M.; Jassal, B.; Stephan, R.; Milacic, M.; Rothfels, K.; Senff-Ribeiro, A.; Griss, J.; Sevilla, C.; Matthews, L.; Gong, C.; et al. The reactome pathway knowledgebase 2022. Nucleic Acids Res 2022, 50 (D1), D687–D692. DOI: 10.1093/nar/gkab1028

(18) Gu, Y.; Zhou, Y.; Ju, S.; Liu, X.; Zhang, Z.; Guo, J.; Gao, J.; Zang, J.; Sun, H.; Chen, Q.; et al. Multi-omics profiling visualizes dynamics of cardiac development and functions. Cell Rep 2022, 41 (13), 111891. DOI: 10.1016/j.celrep.2022.111891

